# mTOR-activating mutations in *RRAGD* cause kidney tubulopathy and cardiomyopathy (KICA) syndrome

**DOI:** 10.1101/2021.03.11.434334

**Authors:** Karl P. Schlingmann, François Jouret, Kuang Shen, Anukrati Nigam, Francisco J. Arjona, Claudia Dafinger, Pascal Houillier, Deborah P. Jones, Felix Kleinerüschkamp, Jun Oh, Nathalie Godefroid, Mehmet Eltan, Tülay Güran, Stéphane Burtey, Marie-Christine Parotte, Jens König, Alina Braun, Caro Bos, Maria Ibars Serra, Holger Rehmann, Fried J.T. Zwartkruis, Kirsten Y. Renkema, Karin Klingel, Eric Schulze-Bahr, Bernhard Schermer, Carsten Bergmann, Janine Altmüller, Holger Thiele, Bodo B. Beck, Karin Dahan, David Sabatini, Max C. Liebau, Rosa Vargas-Poussou, Nine V.A.M. Knoers, Martin Konrad, Jeroen H.F. de Baaij

## Abstract

**Background:** Over the last decaces, advances in genetic techniques have resulted in the identification of rare hereditary disorders of renal magnesium and salt handling. Nevertheless, ±20% of all tubulopathy patients remain without genetic diagnosis. Here, we explore a large multicentric patient cohort with a novel inherited salt-losing tubulopathy, hypomagnesemia and dilated cardiomyopathy (DCM).

**Methods:** Whole exome and genome sequencings were performed with various subsequent functional analyses of identified *RRAGD* variants in vitro.

**Results:** In 8 children from unrelated families with a tubulopathy characterized by hypomagnesemia, hypokalemia, salt-wasting, and nephrocalcinosis, we identified heterozygous missense variants in *RRAGD* that mostly occurred *de novo*. Six of these patients additionally suffered from DCM requiring heart transplantation in 3 of them. An additional dominant variant in *RRAGD* was simultaneously identified in eight members of a large family with a similar renal phenotype. *RRAGD* encodes GTPase RagD mediating amino acid signaling to the mechanistic target of rapamycin complex 1 (mTORC1). RagD expression along the mammalian nephron include the thick ascending limb and the distal convoluted tubule. The identified *RRAGD* variants were shown to induce a constitutive activation of mTOR signaling *in vitro*,

**Conclusions:** Our findings establish a novel disease phenotype combining kidney tubulopathy and cardiomyopathy (KICA) caused by an activation of mTOR signaling suggesting a critical role of Rag GTPase D for renal electrolyte handling and cardiac function.

**SIGNIFICANCE STATEMENT:** Here, we report on heterozygous variants in *RRAGD* in patients with profound hypomagnesemia, renal salt wasting, nephrocalcinosis, and dilated cardiomyopathy. The identified RagD variants induce a constitutive activation of mTOR signaling *in vitro*. These findings not only establish a novel monogenic disorder of the kidney tubule, but demonstrate the essential role of mTOR signaling for distal tubular electrolyte handling and cardiac function.

## INTRODUCTION

The kidney tubule balances fluid, electrolyte and mineral homeostasis. The genetic deciphering of hereditary disturbances of electrolyte reabsorption by the tubule, collectively known as tubulopathies, has greatly improved the understanding of electrolyte transport along the nephron.(1, 2) Over the last decade, we and others have identified critical components of epithelial Mg^2+^ reabsorption pathways in the thick ascending limb of Henle’s loop (TAL) and the distal convoluted tubule (DCT).(2, 3) In the TAL, 50-70% of the filtered Mg^2+^ is reabsorbed paracellularly via the claudin16-19 complex.(4) This paracellular Mg^2+^ transport is dependent on the transcellular salt reabsorption and negatively regulated by basolateral calcium-sensing receptor.(3, 5) In the DCT, Mg^2+^ reabsorption is mediated by transient receptor potential melastatin type 6 and 7 (TRPM6/TRPM7) channels.(6-12)

In our cohort of patients with hypomagnesemia, we noted a subset of patients sharing a tubulopathy characterized by (i) renal salt wasting, (ii) profound hypomagnesemia, and (iii) nephrocalcinosis, combined with a dilated cardiomypathy (DCM). As DCM had not been described in patients with known hereditary forms of hypomagnesemia,(2) we hypothesized that these individuals were affected by a yet undefined disease phenotype. By next-generation sequencing, we identified *RRAGD* variants in eight unrelated individuals, as well as in eight members of a large family with symptomatic hypomagnesemia.

## METHODS

### Subjects

We initially studied a cohort of 6 individuals with a phenotype combining DCM and a complex renal tubular disorder, including hypomagnesemia (median serum magnesium level of 0.42 mmol/L). Our second cohort consisted of 3 additional families in which the index cases presented with hypomagnesemia, hypokalemia, hypercalciuria, and nephrocalcinosis. Clinical and biological data at the time of disease manifestation were collected retrospectively from medical charts. We reevaluated all patients and obtained follow-up biochemical data. The detailed clinical history of each individual is provided as Supplemental Materials. All genetic studies were approved by the local ethics committees in Paris, Münster and Liège, and informed consent was obtained from all subjects and/or their parents.

Serum magnesium, potassium, and chloride levels of individuals with *RRAGD* mutations were statistically compared to non-affected family members. Samples were tested for normal distribution using the Kolmogorov-Smirnov test. For statistical analyses, a one-sided Student t-test in case of normal distribution and a one-sided Mann-Whitney U test in case of non-normal distribution were used.

### Sequencing

Patients F1.1 to F4.1 were subjected to WES (for details see Supplementary Appendix). Exome data were analyzed for shared genes with variants under all modes of inheritance. After discovery of heterozygous *RRAGD* variants in these 4 patients, 2 additional patients with DCM and renal tubulopathy were analyzed by conventional Sanger sequencing (F5.1, F6.1). Then, the analyses were expanded to 25 patients with a similar renal tubular disorder, but without known cardiac phenotype that revealed 2 additional patients (F7.1, F8.1).(40) In short, genomic DNA was fragmented, amplified and sequenced simultaneously by Illumina next generation sequencing technology using an Illumina NextSeq system. DNA samples of members III.1, III.6, and IV.2 family 9 were subjected to WGS on the Illumina X Ten platform (Hartwig Medical Foundation Amsterdam, The Netherlands). Data analysis was performed by mapping paired end reads (2×100 bp) from the NextSeq instrument against the hg19 human reference genome using Burrows Wheeler Aligner with recommended standard settings. Filtering steps excluded common SNPs (> 1%), introns, untranslated regions (UTR), synonymous variants and low-coverage regions (< 5 reads). Overlaps of the remaining variants were generated using VCFMiner to select genes that were present in F1.1-F4.1, which resulted in the identification of one remaining gene, *RRAGD*.(41) Subsequently, individuals F5.1 and F6.1 were subjected to conventional Sanger sequencing of the complete *RRAGD* gene. An extensive description of the sequencing procedure has been provided in the Supplemental Data.

Due to the lack of genetic material, a *RRAGD* mutation could not be confirmed in the affected father of individual F4.1 who died from DCM at 41 years of age. Of note, an older sister of individual F7.1 died in infancy from acute cardiac failure during a pneumonia episode and the autopsy revealed DCM. Careful genetic reevaluation revealed mosaicism (17%) in leucocytes of the healthy mother for the respective *RRAGD* mutation (see Supplemental Data).

### Cell Lysates and Immunoprecipitation

Human embryonic kidney (HEK293T) cells were transfected with Flag-S6K1, RagA and RagD mutants as described previously.(23) To determine the response of mTORC1 to amino acid stimulation or starvation under the control of Rag mutants, two million HEK293T cells were plated onto a 10-cm dish. Twenty-four hours later, the cells were transfected with 2 ng of S6K1 and 100 ng of each Rag wildtype or mutant construct. Thirty-six hours later, amino acid stimulation or starvation was performed for 0.5-1 hour. Cells were rinsed once with ice-cold PBS and lysed with Triton lysis buffer (40 mM Na-HEPES, pH 7.4; 5 mM MgCl2; 100 mM ATP; 10 mM Na4P2O7; 10 mM Na β-glycerol phosphate; 1% v/v Triton; and one tablet of protease inhibitor cocktail per 25 mL of buffer). The lysates were cleared by centrifugation at 15,000 rpm at 4°C in a microcentrifuge for 10 minutes. For immunoprecipitations, the FLAGM2 beads were pre-equilibrated in Triton lysis buffer. 30 µL of a 50/50 slurry of the FLAG-M2 affinity beads were then added to cleared lysates and incubated at 4°C for two hours. Following immunoprecipitation, the beads were washed one time with Triton lysis buffer and 3 times with Triton lysis buffer supplemented with 500 mM NaCl. Immunoprecipitated proteins were denatured by the addition of 50 µL of 2.5 x SDS buffer, resolved by SDS-PAGE, and analyzed by immunoblotting.

### Sample preparation for interactome analysis

FlpIn-IMCD (inner medullary collecting duct) cells stably expressing GFP-GFP, GFP-Rragd-wt or GFP-Rragd-Ser75Leu fusion proteins were generated as described previously.(42) Lysates were cleared by centrifugation, followed by incubation with GFP µMACS magnetic beads (Miltenyi) for 1 h. Then, lysates were loaded on a µMACS column and precipitated proteins were reduced, alkylated and on-column digestion was performed over-night as previously described. Eluates were acidified the next day using 1-2% formic acid and stage-tip clean-up was performed as previously described and samples were dried using a vacuum centrifuge. Peptides were resuspended in 0.1% FA and were separated using a 1h gradient on an nLC coupled to a Q Exactive Plus tandem mass spectrometer (Thermo scientific) or an LTQ orbitrap XL mass spectrometer. Experimental details are provided in the Supplemental Data.

### Expression analysis

Total RNA was extracted from issues of C57BL/6 mice and isolated distal convoluted tubule (DCT) cells of parvalbumin-GFP mice using Trizol Reagent (Thermo Fisher Scientific, Waltham, MA, USA) according to manufacturer’s protocol. To remove genomic DNA, the isolated RNA was subsequently subjected to DNase (Promega, Madison, WI, USA) treatment. RNA was reverse transcribed using Moloney murine leukemia virus reverse transcriptase (Thermo Fisher Scientific) for 1 h at 37°C. Ras-related GTP binding D (Rragd) (forward, CACCTGAGCTTTTACCTGA; reverse, TCAGCAGATTCTCCAGCGTC) gene expression levels were quantified by SYBR-Green (Bio-Rad) on a CFX96 Real-Time PCR Detection System (Bio-Rad) and normalized for glyceraldehyde 3-phosphate dehydrogenase (Gapdh) (forward TAACATCAAATGGGGTGAGG; reverse, GGTTCACACCCATCACAAAC) expression levels.

Immunohistochemistry was performed on kidney tissues of C57BL/6 mice using primary antibodies against RagD, AQP2, NCC, and uromodulin as described previously (43). Paraffin-ebedded kidney sections were subjected to deparaffinization and rehydration, followed by permeabilization in PBS-Triton 0.3% (v/v) (Sigma). Sections were incubated overnight at 4°C with the primary antibodies sheep anti-NCC (1:400) (Sheep S965B, MRC PPU, Dundee), sheep anti-Tamm Horsfall (1:200) (MBS220487, Bio-Trend) and guinea pig anti-AQP2 (1:100) (kindly provided by dr. Peter Deen) and subsequently incubated overnight at 4°C with the primary antibody rabbit anti-RRAGD (1:2,000) (NBP2-32106, Novus Biologicals). Experimental details are provided in the Supplemental Data.

## RESULTS

### Patients

We initially identified eight unrelated individuals with hypomagnesemia and a complex renal tubulopathy phenotype (Table 1), accompanied by muscle spasms and seizures in three individuals. Inappropriately high fractional excretion rates of magnesium indicated renal magnesium wasting (FE-Mg, Table 1). In individual F2.1, a magnesium loading test was performed, which disclosed a decreased renal tubular threshold for magnesium reabsorption (Figure 1A). Additional renal findings included polyuria, hypokalemia, hypochloremia, a tendency towards metabolic alkalosis, hypercalciuria, and nephrocalcinosis pointing to a defect in the TAL (Table 1, Figure 1B-E). Of note, the prenatal course of individuals F3.1 and F4.1 was complicated by polyhydramnios, as observed in antenatal Bartter syndrome.(13). All family members of these probands were clinically unaffected. In six patients (F1-F6), the tubulopathy was accompanied by DCM, resulting in terminal heart failure and requiring heart transplantation in individuals F1.1 to F3.1 (Figure 1F).

**Table 1.**
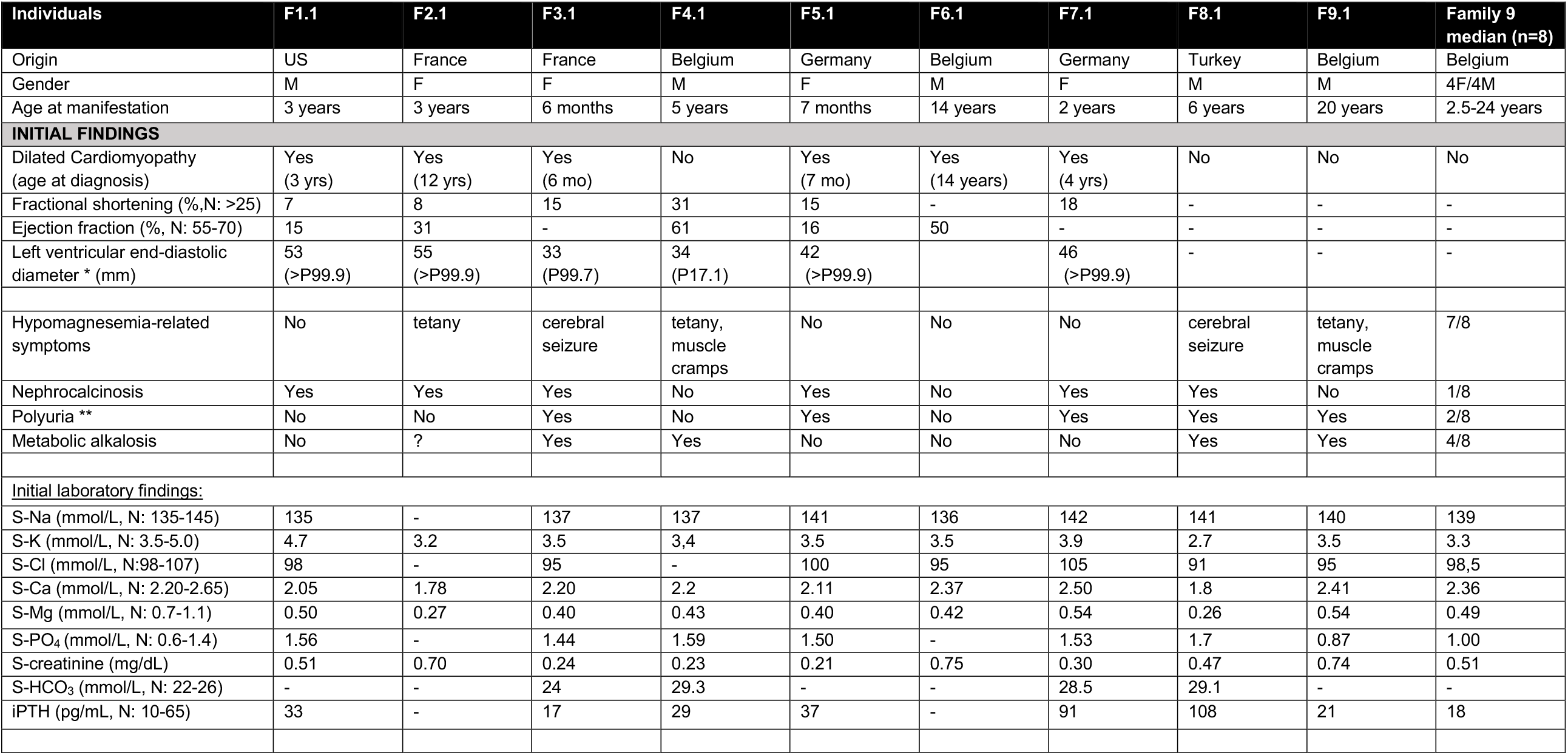
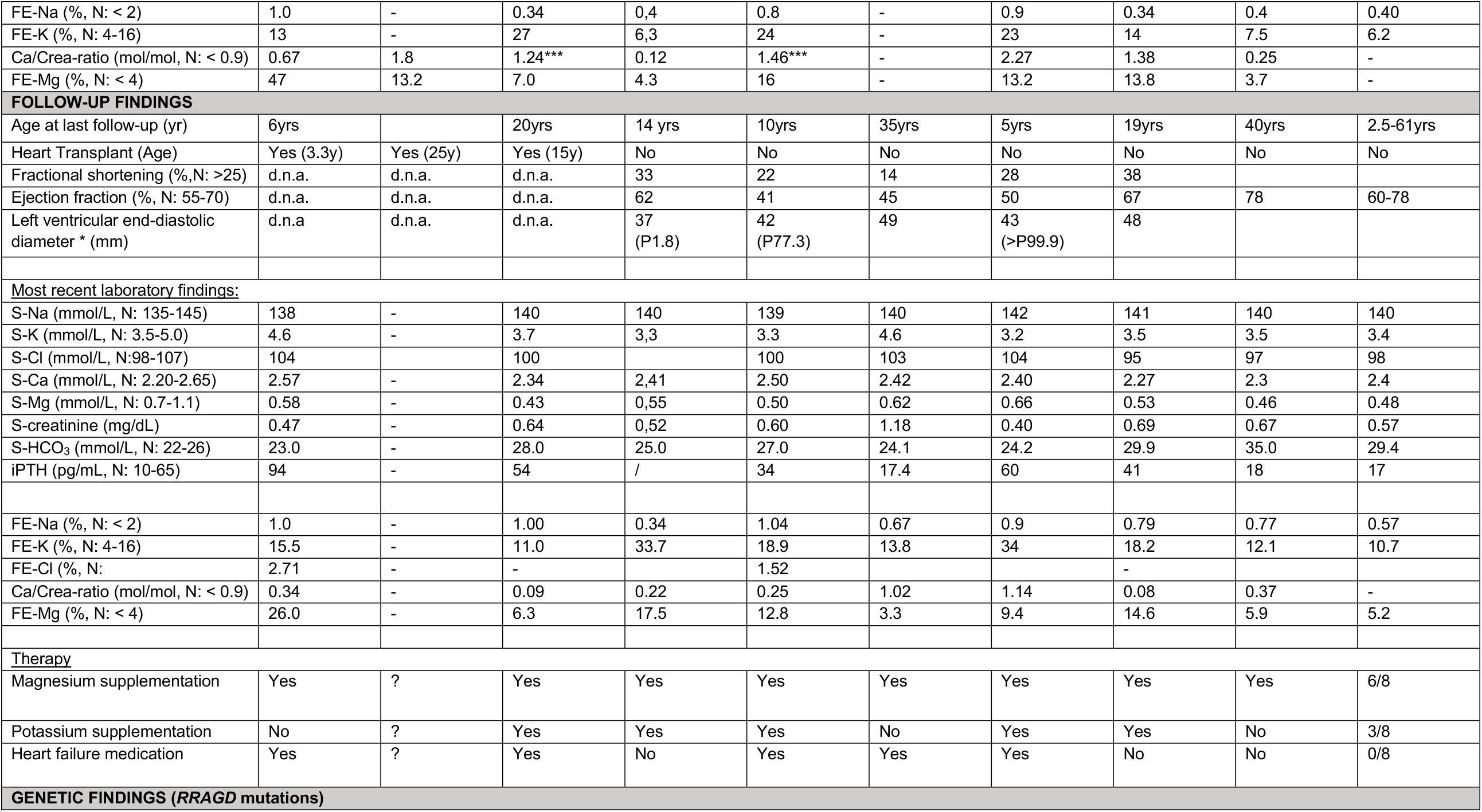
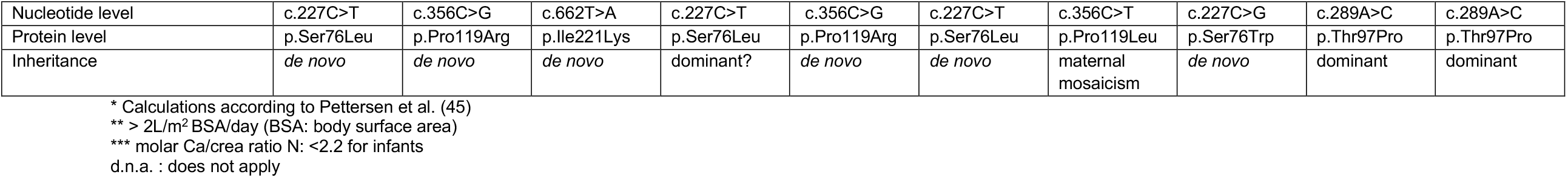
Clinical and laboratory characteristics of patients with *RRAGD* mutations

**Figure 1.**
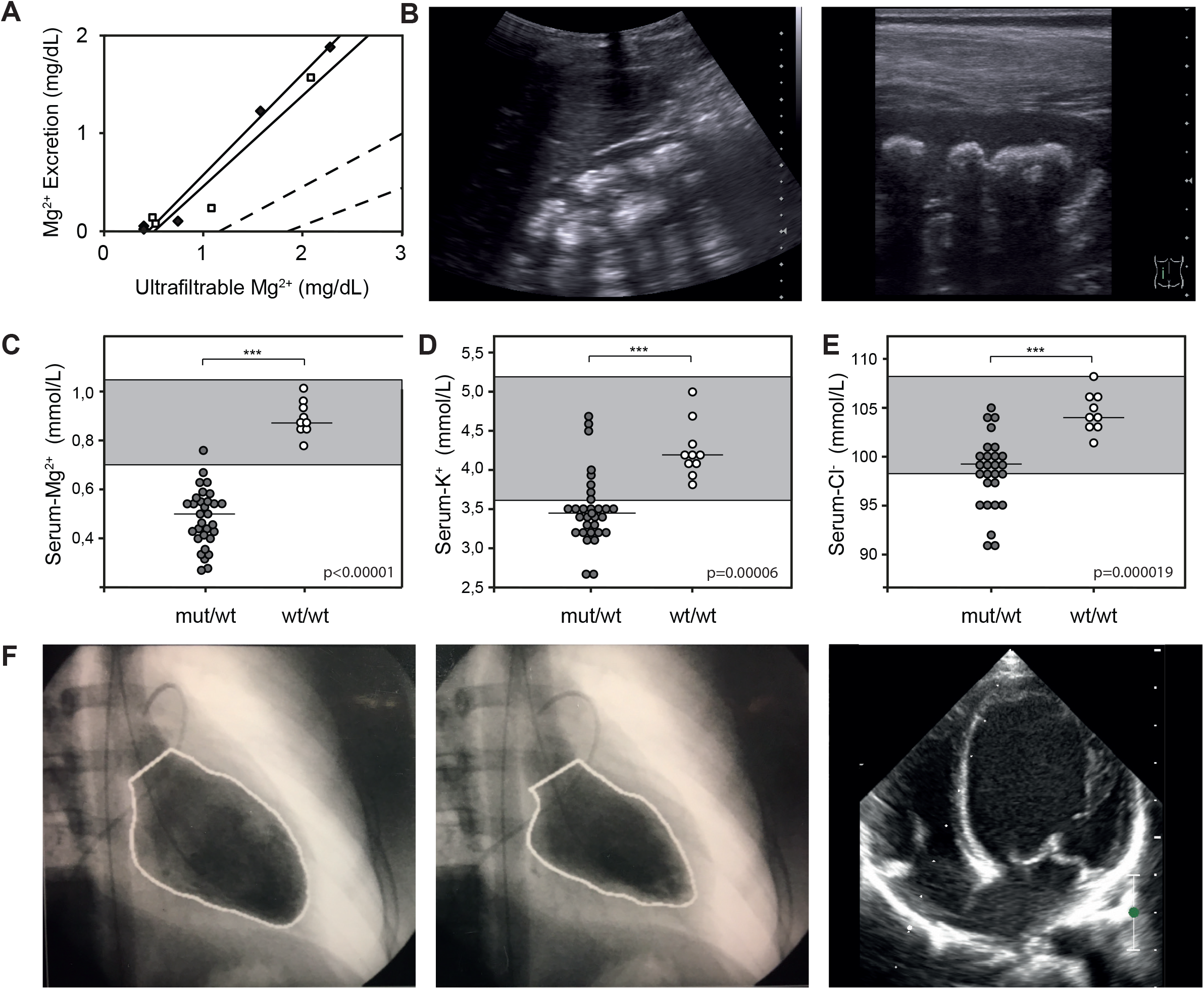
Clinical and genetic findings in 9 families with cardiomyopathy and renal tubulopathy. (**A**) Magnesium loading test in individual F2.1. Urinary magnesium excretion is plotted against the ultrafiltrable fraction of serum magnesium before and during parenteral infusion of magnesium at increasing concentrations (two separate tests). The dashed lines represent the normal range. (**B**) Renal ultrasound of individual F7.1 showing severe medullary nephrocalcinosis (Grade III according to Hoyer)(44). (**C-E**) Comparison of serum magnesium, potassium and chloride levels of individuals with *RRAGD* mutations (grey circles) and of unaffected family members (white circles)(initial as well as follow-up values included). Horizontal bars represent medians, reference ranges for serum electrolytes are indicated by grey boxes. Next to significant hypomagnesemia, individuals with *RRAGD* mutations exhibited significantly lower serum potassium and serum chloride levels reflecting renal salt wasting (one-sided Student’s t-test (magnesium, chloride) and Mann Whitney U-test (potassium); *** = p<0.001). (**F**) Left ventriculography (performed at the age of 12 years) of individual F2.1 who presented with cardiomyopathy. Pictures show left ventricular volumes during diastole (left) and systole (middle) demonstrating a reduced ejection fraction of 40% (normal 55-70%). Initial echocardiogram of individual F7.1 performed after genetic diagnosis (apical 4-chamber view) showing the dilated left ventricle.

In parallel of these eight unrelated patients, we analyzed a large family (F9) with eight affected members presenting with profound and symptomatic hypomagnesemia resistant to adequate oral magnesium supplements (pedigree and clinical details in Supplementary Figure 1 &Supplementary Table 1, respectively). The index case presented in early adulthood with muscle cramps and tetany. In addition, affected individuals exhibited hypokalemia, metabolic alkalosis and an activation of the renin-angiotensin-aldosterone system (RAAS) together with normal or low blood pressure suggesting the diagnosis of Gitelman syndrome (Table 1, Figure 1C). None of these patients showed cardiac abnormalities.

Serum potassium and chloride levels were significantly lower in all affected individuals (n=16) compared with those of available non-affected family members (n=8) (Figure 1D-E), which, together with metabolic alkalosis and RAAS activation observed in family 9, was highly indicative of renal salt wasting.

### Identification of *RRAGD* mutations

After exclusion of mutations in known genes for hypomagnesemia and salt wasting, we performed whole-exome sequencing (WES) in individuals F1.1 to F4.1, which revealed *RRAGD* as a single common mutated gene. The identified *RRAGD* variants were all present in heterozygous state and mostly occurred *de novo*. Subsequently, additional heterozygous *RRAGD* variants were identified in four sporadic individuals F5.1 to 8.1. Family F9 was analyzed separately under the assumption of a dominant trait. Whole genome sequencing (WGS) revealed a heterozygous *RRAGD* variant (p.Thr97Pro) that co-segregated with the disease phenotype.

*RRAGD* encodes a small Rag guanosine triphosphatase (GTPase), which is a member of the Ras family of GTP-binding proteins.(14) All RagD mutations affect highly conserved amino acid residues in GTP-binding domains conserved in small GTPases (G-domains, G1 to G5, Figure 2A-D, Supplementary Figure 2).(15)

**Figure 2.**
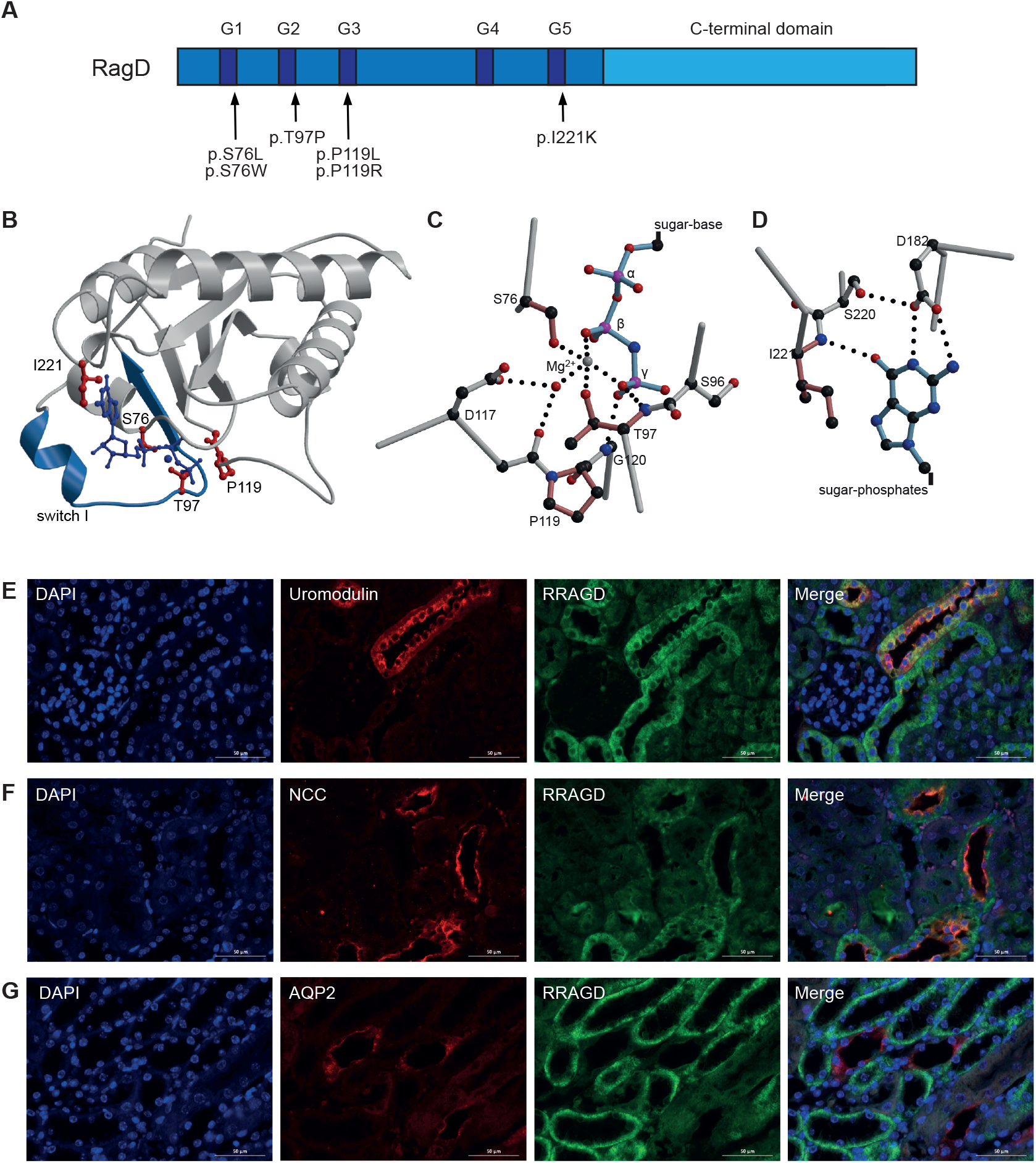
Localization of mutations in the RagD protein. (**A**) Domain organization of RagD with GTP binding motifs in dark blue (G1-G5). Mutations are indicated with an arrow. (**B**) Crystal structure of RagD in complex with a GTP analogue (GppNHp, pdb entry 2q3f). Mutated residues are shown in red. GppNHp and the coordinated Mg^2+^ ion are depicted in dark blue. (**C-D**) Detailed view of the phosphate moiety of the nucleotide (C) and the nucleotide base (D). Affected residues are colored in light red. Dotted lines indicate hydrogen bonds. (**E-G**) Double immunofluorescence staining of mouse kidney (B-D) sections for RagD in green and uromodulin (B, red), NCC (C, red) or aquaporin (D, red). Bars represent 20µm.

RagD showed a ubiquitous expression pattern with significant transcript levels in heart and kidney (Supplementary Figure 3). Compared to well-established markers of distal tubular segments, RagD co-localized with uromodulin (UMOD) and Na^+^-Cl^-^-cotransporter (NCC) in the TAL and DCT nephron segments, respectively (Figure 2E-G).

### RagD mutations activate the mTOR pathway

All RagD mutants showed increased binding to Raptor (regulatory associated protein of mTOR) and mTOR, the essential components of mTORC1 (Figure 3A) after exogenous expression in HEK293T cells. All human RagD mutants resulted in increased S6K1 phosphorylation in the presence of nutrients, indicative of active mTORC1 signaling. Of note, some mutants increased S6K1 phosphorylation even during starvation (Figure 3B).

**Figure 3.**
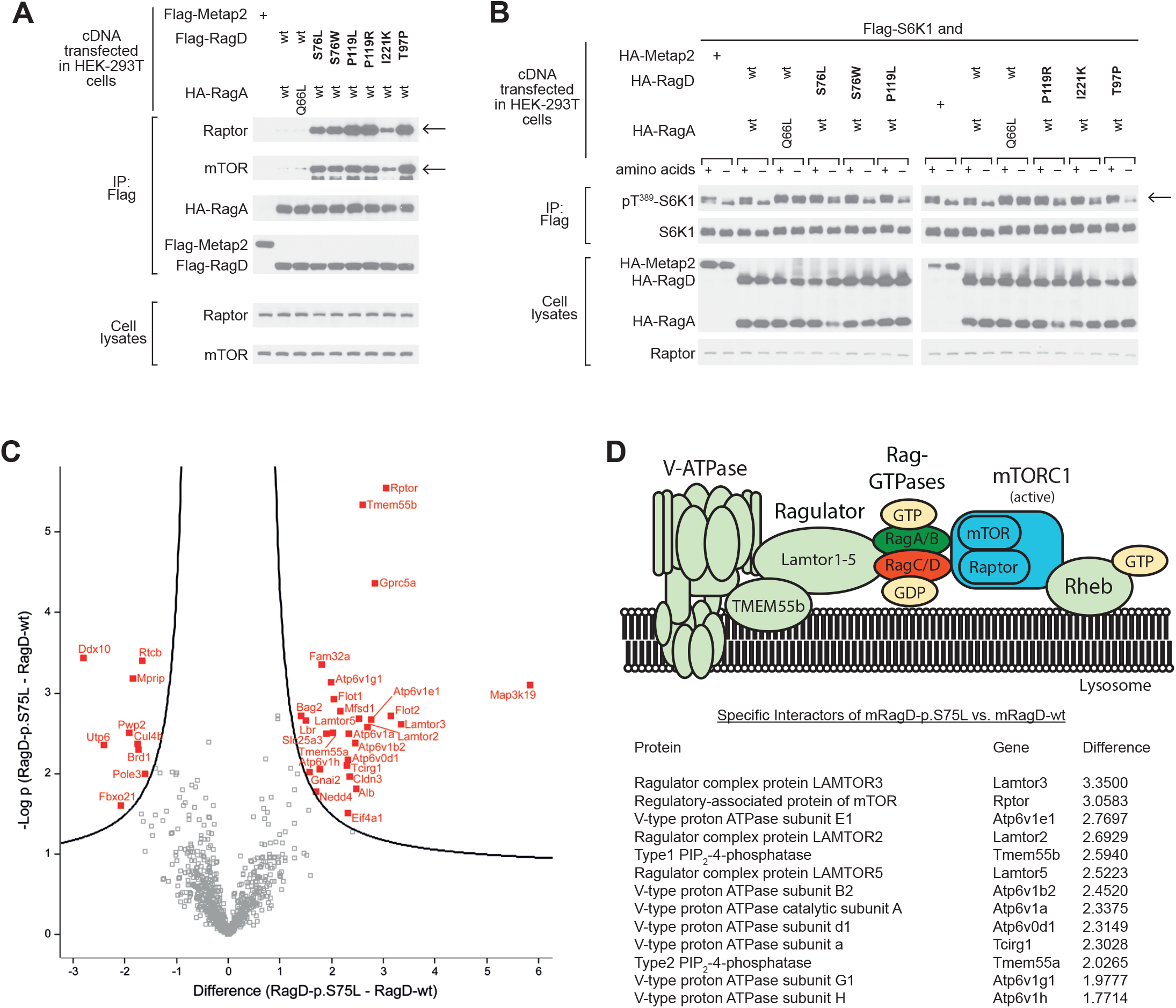
*RRAGD* mutations increase mTOR signaling *in vitro*. (**A**) Immunoblot analysis demonstrating that RagD mutations (bold) increase Raptor and mTOR binding, indicated by arrows (top rows). RagA and RagD blots indicate equal (co-) precipitation of the Rag dimer (middle rows). Transfected HEK293T cells contained equal amounts of Raptor and mTOR as detected in the cell lysate fraction (bottom rows). (**B**) mTORC1 signaling in response to amino acids and starvation assessed by phosphorylation of its substrate S6 kinase (S6K1) at Threonine 389 (Thr^389^), indicated by an arrow. Compared to wt RagD, all RagD mutants caused an increased S6K1 phosphorylation in the presence of amino acids. The expression of transfected RagA, RagD and S6K1, as well as of endogenous Raptor, was confirmed by immunoblotting (bottom rows). Constitutively active RagA-p.Q66L served as a positive control, (25) Metap-2 as a negative control for non-specific flag-binding. (**C**) Volcano plot of the interactome of murine RagD-p.S75L vs. RagD-wt. The significance (p-value) of changes between RagD-p.S75L and RagD-wt (-log10 scale) is plotted against the difference (fold change) between RagD-p.S75L and RagD-wt (log2 scale). Dots beyond the curved line represent proteins with significantly increased abundance (FDR 0.1, s0=0.5) in RagD-p.S75L samples (right) or RagD-wt samples (left). (**D**) Model of the V-ATPase-Ragulator-Rag-GTPase-mTORC1 complex and table listing the identified components from panel C that showed significantly increased binding to RagD-p.S75L in log2-fold change.

To further evaluate the effect of human RagD-p.Ser76Leu on mTORC1 complexes, we performed an interactome analysis by quantitative proteomics after stable genomic integration in a murine tubular epithelial cell line (Figure 3C). Here, mutant RagD-p.Ser75Leu showed a significantly increased co-precipitation of multiple proteins of the mTORC1 signaling complex (Figure 3D), includingRaptor, members of the Lamtor protein family (Lamtor3, Lamtor5) and different subunits of lysosomal vacuolar-type H^+^-ATPase (V-ATPase, Atp6v-proteins), (Figure 3C-D, Supplementary Figure 4).

## DISCUSSION

The present study describes a previously unrecognized hereditary disease phenotype combining an electrolyte-losing tubulopathy and DCM that is caused by heterozygous missense variants in the *RRAGD* gene encoding Rag GTPase D. The identified *RRAGD* variants lead to increased mTOR signaling. We propose to call this novel inherited disease: “the KICA (**Ki**dney tubulopathy and **Ca**rdiomyopathy) syndrome”.

The cardinal symptom of individuals with *RRAGD* mutations is a severe and substitution-resistant hypomagnesemia, resulting in muscle cramps, tetany, and cerebral seizures. Renal magnesium wasting was associated with hypercalciuria and nephrocalcinosis in a subset of individuals, a phenotype highly reminiscent of FHHNC (Familial Hypomagnesemia with Hypercalciuria and Nephrocalcinosis; OMIM #248250, #248190)(2, 3). In FHHNC, mutations in two tight junction proteins, claudin-16 and claudin-19, cause a defective paracellular reabsorption of calcium and magnesium in the TAL.(16-18) Immunostaining experiments indeed showed RagD expression in the TAL, suggesting that RagD variants may impair Ca^2+^ and Mg^2+^ reabsorption in this tubular segment.

In a subset of patients, a renal phenotype with salt wasting and hypokalemic alkalosis prevailed resembling Bartter syndrome. The prenatal finding of polyhydramnios in three individuals supported this assumption. Unfortunately, a systematic evaluation of the RAAS in our entire cohort was hampered by early cardiac insufficiency and treatment with diuretics and ACE inhibitors in most individuals. However, a marked activation of the RAAS was also noted in individual F2.1 years before cardiac manifestation supporting the suspicion of a mixed FHHNC/salt wasting phenotype. On the other hand, hypercalciuria and nephrocalcinosis lacked in affected members of family 9. This phenotypic variability might be explained by a less pronounced activation of mTOR signaling by the RagD-p.T97P mutant, but could also reflect a primary dysfunction of the DCT with a Gitelman syndrome-like phenotype rather than a TAL defect. Similar observations have been made in Bartter syndrome type III due to *CLCNKB* defects, in which individuals might present with a phenotype ranging from antenatal Bartter syndrome to Gitelman syndrome.(19, 20) Future research will be required to elucidate the exact targets of increased mTOR signaling in distal tubular segments.

Interestingly, the phenotype of our cohort is reminiscent of 2 individuals from a previously described Finnish family with unresolved renal tubular disease and cardiomyopathy.(21, 22) The clinical picture of these individuals is highly suggestive of KICA syndrome. Unfortunately, the cardiac phenotype resulted in an early fatal issue in this family, which hampers any further genetic investigation.

RRAGD encodes a small Rag guanosine triphosphatase (GTPase), which is an essential component of the nutrient-sensing pathway that activates mTOR signaling (14). mTOR serves as the main nutrient sensor of the cell coordinating signals from extracellular growth factors and intracellular nutrient availability, such as amino acids.(23) (24) Upon amino acid stimulation, Rag GTPases target the mTOR complex1 (mTORC1) to the lysosome where its kinase is activated.(23) Indeed, our interaction studies demonstrate increased interaction of RagD-S76L to components of the mTOR signaling pathway and the Ragulator complex (Figure 3). The human *RRAGD* mutations discovered in the present study affect residues involved in the binding of guanosine nucleotides (Figure 2, Supplementary Figure 2). Previously, engineered mutations that interfere with nucleotide binding have been shown to critically affect Rag signaling and mTORC1 activation.(23, 25) In line with this, the variants discovered here lead to an increased activation of mTORC1, a subset even under amino acid starvation (Figure 3).

Data on the role of mTOR signaling in tubular physiology is still scarce. Previous studies mainly focused on the tubular effects of decreased mTORC1 signaling.(26, 27) Inactivation of mTOR signaling along the renal tubule was shown to lead to a Fanconi-like syndrome as well as a urinary concentrating defect attributed to impaired distal tubular transport processes.(28) Remarkably, hypomagnesemia and hypokalemia have been observed in rats after sirolimus treatment that were attributed to reduced expression of NKCC2 (SLC12A1) in the TAL.(29) Moreover, inhibition of mTOR signaling also resulted in changes in TRPM6 expression in the DCT.(30) Conversely, mTORC1 activation by knock-out of *Tsc1* in renal tubular cells resulted in cyst formation resembling polycystic kidney disease (PKD).(31, 32) Our findings now underline the indispensable role of mTORC1 signaling for tubular electrolyte handling that appears to be particularly sensitive to both inactivation and hyperactivity of the mTOR pathway.

The finding of DCM in a subset of our patients also substantiates a critical role of Rag GTPases for cardiac function. Previously, inactivation of Rag GTPases RagA and RagB was shown to result in hypertrophic cardiomyopathy in mice.(33) Moreover, a *de novo* mutation in RagC (p.Ser75Tyr) leading to increased mTOR activity has been described in a single individual with DCM.(34) Recent studies in clinical and experimental models of DCM have implicated mTOR hyperactivation as a major cause of DCM.(35, 36) In line, inhibition of mTOR by rapamycin has been demonstrated to successfully improve cardiac function in animal models of DCM.(37-39) Of note, not all identified *RRAGD* mutations overactivated mTORC1 signaling equally. Our experiments indicate that the p.Thr97Pro results in less S6K1 activation (Figure 3), which may explain why individuals carrying this mutation do not suffer from DCM even at adult age.

Taken together, the discovery of *RRAGD* mutations establishes a novel hereditary tubulopathy linked to dysregulation of the mTOR pathway. The diagnosis of KICA syndrome should be considered in individuals presenting either with early-onset DCM and/or hypomagnesemia of renal origin. Future research will hopefully help to enlighten the exact role of mTOR for distal tubular transport processes and renal magnesium handling in particular.

## ACKNOWLEDGEMENTS

The authors are grateful to the patients for their participation in this study. We thank Edith Peters, Femke Latta, Valentina Carotti, Ruud Tilleman, Matthijs Snelders, Marie Neuville, and Ruth Herzog for their excellent technical support and Tanja Seidel for thoughtful patient care. This work was supported by grants from the Netherlands Organization for Scientific Research (NWO VENI 016.186.012), the Dutch Diabetes Research Foundation (2017.81.014), the Dutch Kidney Foundation (15OKG18 to KYR), and the EURenOmics project from the European Union seventh Framework Program (FP7/2007-2013, no. 305608). FJ is a Fellow of the Fonds National de la Recherche Scientifique (FNRS) in Belgium.

## SUPPLEMENTAL DATA – TABLE OF CONTENTS

### Text

1. Complete methods
2. Detailed clinical description of affected individuals
3. Detailed description of the structural consequences of the mutations
4. References

### Tables

1. Biochemical parameters F9

### Figures

1. Pedigree F9
2. RagD mutations affect conserved residues
3. Tissue expression of RagD
4. Effects RagD mutations on its interactome

