## Supplement for "mTOR-activating mutations in *RRAGD* cause kidney tubulopathy and cardiomyopathy (KICA) syndrome"

**SUPPLEMENTAL DATA**

**Text**

1. Complete methods

2. Detailed clinical description of affected individuals

3. Detailed description of the structural consequences of the mutations

4. References

**Tables**

1. Biochemical parameters F9

**Figures**

1. Pedigree F9

2. RagD mutations affect conserved residues

3. Tissue expression of RagD

4. Effects RagD mutations on its interactome

### COMPLETE METHODS

#### Whole Exome Sequencing

Genomic DNA was extracted from EDTA blood according to standard procedures. Individuals F1.1, F2.1, F3.1, F4.1 F7.1 and F8.1 were subjected to whole exome sequencing as described previously.[1] In short, we used a customized sequence capture library that targeted exons and additional 35 bp of flanking intronic sequence. Genomic DNA was fragmented, and the coding exons as well as the corresponding exon-intron boundaries were enriched using the Roche/NimbleGen sequence capture approach (NimbleGen, Madison, Wisconsin, USA). They were then amplified and sequenced simultaneously by Illumina next generation sequencing (NGS) technology using an Illumina NextSeq system. NGS data analysis was performed by mapping paired end reads (2×100 bp) from the NextSeq instrument against the hg19 human reference genome using BWA with recommended standard settings. Mapped reads were preprocessed with SAM tools and duplicate reads were marked by Picard. Finally, GATK was applied for local realignment and base quality score recalibration of mapped reads. Filtering steps excluded common SNPs (> 1%), introns, untranslated regions (UTR), synonymous variants and low-coverage regions (< 5 reads). Overlaps of the remaining variants were generated using VCFMiner to select genes that were present in F1.1-F4.1 using the following settings: non-synonymous variants in exons and canonical splice sites, with a minor allele frequency from Exac, and in house data of less than 1%. This analysis resulted in the identification of 1 remaining gene *RRAGD*.[2] Subsequently, patients F5.1 and F6.1 were subjected to Sanger sequencing of the complete *RRAGD* gene.

#### Whole Genome Sequencing (WGS) in family 9

Genomic DNA was extracted from 10 ml EDTA blood according to standard and automated procedures. DNA samples of family 9, members III.1, III.6, and IV.2, were subjected to WGS on the Illumina X Ten platform (Hartwig Medical Foundation Amsterdam, The Netherlands). WGS reads were processed by following the Genome Analysis ToolKit (GATK) best practices guidelines[1] where FASTQC was applied for quality control of the raw data. Sequencing reads were mapped onto the human genome (GRCh37/hg19) using Burrows Wheeler Aligner (v.0.7.5a). As part of GATK best practices, duplicate reads were removed with Picard (v.1.119), VariantRecalibrator was used for recalibration of base quality scores, and single nucleotide variants (SNVs) and indels were called using HaplotypeCaller (v.3.2-2). Next, copy number variants were called using FreeC (v.10.4) and structural variants were called using Delly (v.0.7.2) and Manta (v.1.0.3). The variants called were further prioritized and classified using Agilent Cartagenia software (v.4.3.5) based on the American College of Medical Genetics (ACMG) guidelines on SNV interpretation[2], population frequency (Minor Allele Frequency; MAF < 0.0001) acquired from the 1000 Genomes Phase 1 & 3, Exome Aggregation Consortium (ExAC) and the Genome of The Netherlands (GoNL v.5), location of the variants (exonic, 5' untranslated region, 3' untranslated region, intronic regions limited to 20 base pairs upstream and downstream of exons, and non-coding RNA exonic regions), variant quality (read depth ≥ 10 and filter status = PASS), conservation scores, and *in silico* predicted effects on gene products and functions (SIFT ≤ 0.5, PolyPhen-2 HumDiv = possibly or probably damaging, PolyPhen-2 HumVar = possibly or probably damaging, Mutation Taster = disease-causing, and FATHMM ≤ 0 ). Final fine-tuning filtering steps involved overlap of variants in family members III.1, III.6, IV.2 and determination of the novelty of overlapping variants identified, (MAF = 0; using Genome Aggregation Database; https://gnomad.broadinstitute.org/), exploring literature and databases on known biological information from sources such as Online Mendelian Inheritance in Man (https://www.omim.org/) and The Human Protein Atlas (https://www.proteinatlas.org). Three overlapping heterozygous variants among the three patients were confirmed and checked for co-segregation in the family by Sanger sequencing (Macrogen®, Amsterdam, The Netherlands).

**Next generation sequencing for the detection of maternal mosaicism in family 7**

We used a customized sequence capture library [Lu et al., 2017] that targeted exons and additional 35 bp of flanking intronic sequence. Genomic DNA was fragmented, and the coding exons as well as the corresponding exon-intron boundaries were enriched using the Roche/NimbleGen sequence capture approach (NimbleGen, Madison, Wisconsin, USA). They were then amplified and sequenced simultaneously by Illumina next generation sequencing (NGS) technology using an Illumina NextSeq system. NGS data analysis was performed by mapping paired end reads (2×100 bp) from the NextSeq instrument against the hg19 human reference genome using Burrows Wheeler Aligner with recommended standard settings. Mapped reads were preprocessed with SAM (Sequence Alignment/Map) tools and duplicate reads were marked by Picard. Finally, the Genome Analysis ToolKit (GATK) was applied for local realignment and base quality score recalibration of mapped reads. JSI Medical Systems software (version 4.1.2, SeqNext module) was used for visualization.

**Cell culture and transfection**

HEK293T cells were obtained from ATCC and maintained in an incubator set at 37°C and 5% $CO_2$. HEK293T cells were cultured in DMEM with 10% (v/v) IFS supplemented with 2 mM glutamine, penicillin (100 IU/ml), and streptomycin (100 mg/ml). For co-transfection experiments, 2,000,000 HEK-293T cells were plated in 10 cm culture dishes. Twenty-four hours later, cells were transfected using Xtremegene 9 transfection reagent with the pRK5-based cDNA expression plasmids indicated in the figures.

For stable expression of a GFP-GFP, GFP-RagD-wt or GFP-RagD-p.Ser75Leu fusion protein FlpIn IMCD cells were transfected with the pgLAP3 vector containing murine *Rragd* (WT or Ser75Leu) or GFP CDS, respectively, and Flp-recombinase expression vector (pOG44, Invitrogen) using Lipofectamine 2000 (Invitrogen). Stably integrated cells were selected by Hygromycin B (Invivogen) treatment.

**Cell Lysates and Immunoprecipitation**

To determine the response of mTORC1 to amino acid stimulation or starvation under the control of Rag mutants, two million HEK293T cells were plated onto a 10-cm dish. Twenty-four hours later, the cells were transfected with 2 ng of S6K1 and 100 ng of each Rag wildtype or mutant construct. Thirty-six hours later, amino acid stimulation or starvation was performed for 0.5-1 hour. Cells were rinsed once with ice-cold PBS and lysed with Triton lysis buffer (40 mM Na-HEPES, pH 7.4; 5 mM $MgCl_2$; 100 mM ATP; 10 mM $Na_4P_2O_7$; 10 mM Na $\beta$-glycerol phosphate; 1% v/v Triton; and one tablet of protease inhibitor cocktail per 25 mL of buffer). The lysates were cleared by centrifugation at 15,000 rpm at 4°C in a microcentrifuge for 10 minutes. For immunoprecipitations, the FLAGM2 beads were pre-equilibrated in Triton lysis buffer. 30 μL of a 50/50 slurry of the FLAG-M2 affinity beads were then added to cleared lysates and incubated at 4°C for two hours. Following immunoprecipitation, the beads were washed one time with Triton lysis buffer and 3 times with Triton lysis buffer supplemented with 500 mM NaCl. Immunoprecipitated proteins were denatured by the addition of 50 μL of 2.5 x SDS buffer, resolved by SDS-PAGE, and analyzed by immunoblotting.

**Sample preparation for interactome analysis**

FlpIn IMCD cells expressing a single copy of GFP-GFP, GFP-RagD-wt or GFP-RagD-Ser75Leu fusion proteins were lysed in lysis buffer (1% Triton X-100, 20 mM Tris pH 7.5, 25 mM NaCl, 50 mM NaF, 15 mM Na4P2O7, 1 mM EDTA, 1xPIM, and 5 mM Na3VO4) and by sonication (30% amplitude, 9 s, 0.1 s sonic, 0.9 s pause). Lysates were cleared by centrifugation (17,000 g, 15 min, 4°C), followed by incubation with GFP μMACS magnetic beads (Miltenyi) for 1 h. Then, lysates were loaded on a μMACS column and precipitated proteins were reduced, alkylated and on-column digestion was performed over-night as previously described.[3] Eluates were acidified the next day using 1-2% formic acid and stage-tip clean-up was performed as previously described and samples were dried using a vacuum centrifuge.[4]

**nLC-MS/MS**

Peptides were resuspended in 0.1% FA and were separated using a 1h gradient on an nLC coupled to a Q Exactive Plus tandem mass spectrometer (Thermo scientific) or an LTQ orbitrap XL mass spectrometer. Separation was carried out on an in-house packed 50 cm column with 1.7 µm C18 beads (Dr Maisch GmbH). The gradient consisted of two buffers, Buffer A: 0.1% formic acid and B: 80% acetonitrile, 0.1% formic acid was used. Linear gradients from 7 to 38% B in 60 min were used for separation with a following increase to 80% B for 5 min and a re-equilibration to 5% B. Peptides were then sprayed into a tandem mass spectrometer, the Q Exactive plus (60). The resolution for MS1 spectra was 70 000 (mass range, 200–1200 mz−1). MS1 spectra were acquired using 1E6 as an AGC target. MS/MS spectra of the top 10 most intense peaks were obtained by higher-energy collisional dissociation fragmentation. Resolution for MS/MS spectra was 35 000 at 200 mz−1, AGC target was 5E5, max. injection time was 120 ms.

**Bioinformatic analysis**

RAW files generated by the QExactive tandem mass spectrometer were processed using MaxQuant, v.1.4.1.2 with default settings. Fixed modifications were carbamidomethylation on cysteins. Variable modification was Methionine oxidation. PSM, and site and protein false discovery rate (FDR) was 0.01. Minimal peptide length was 7. Match between run option and label-free quantification option was enabled. Data was searched against a mouse uniprot database in fasta format from the Mus musculus, reference proteome downloaded from uniprot.org on 23rd of February 2014. MaxQuant output (proteingroups.txt file) was uploaded in Perseus, v. 1.6.7.0. Contaminants and reverse hits were removed. Determination of significant interactors was performed using a two-tailed t-test and FDR and s0 cutoff values indicated in the figure legends. Stoichiometry analysis was performed using intensity based absolute quantification (iBAQ) values.

**Immunohistochemistry**

Mouse kidney tissue was fixed in 10% (v/v) neutral-buffered formalin (Klinipath BV) in PBS for >24 h. Samples were dehydrated through alcohol (Klinipath BV), embedded in paraffin (39602004, Leica biosystems), and cut into 4-μm sections. For immunohistochemistry of the kidney sections, deparaffinization and rehydration were performed, followed by permeabilization in PBS-Triton 0.3% (v/v) (Sigma). Sections were incubated overnight at 4°C with the primary antibodies sheep anti-NCC (1:400) (Sheep S965B, MRC PPU, Dundee), sheep anti-Tamm Horsfall (1:200) (MBS220487, Bio-Trend) and guinea pig anti-AQP2 (1:100) (kindly provided by dr. Peter Deen). Subsequently, sections were incubated for 2 h at room temperature with Alexa 594 conjugated secondary antibodies (1:300) (Thermo Fisher Scientific, Molecular Probes). For co-staining with RRAGD, sections were then blocked with 0.3% (v/v) $H_2O_2$ (Denteck) and Avidin/Biotin (SP-2001, Vector Labs) and subsequently incubated overnight at 4°C with the primary antibody rabbit anti-RRAGD (1:2,000) (NBP2-32106, Novus Biologicals). For detection of RRAGD, sections were incubated for 2 hours at room temperature with a swine anti rabbit Biotin SP-conjugated secondary antibody (1:2,000) (4050-08, Southern Biotech), followed by incubation with Streptavidin-HRP (1:100) and fluorescein tyramide (1:50) according to the manufacturers protocol (NEL741001KT, Perkin Elmer). Slides were mounted with Fluoromount-G with DAPI (0100-20, Southern Biotech).

### DETAILED CLINICAL DESCRIPTION

#### Family 1

The boy was born at 38 weeks of gestation after normal pregnancy (normal amniotic fluid) by vaginal delivery. He was reportedly healthy during the first years of life receiving breast feeding for eight months followed by infant formula. There was no indication of vomiting, diarrhea, polyuria or dehydration. At 18 months of age he was admitted to a local hospital for a purple rash on his face and legs, and Henoch-Schonlein Purpura was diagnosed.

At 6 years of age, he presented to a tertiary hospital with a history of decreased oral intake, vomiting, abdominal pain and fatigue. He was found to have severe dilated cardiomyopathy and cardiac insufficiency that did not stabilize under intensive medical therapy and, 7 days after admission, necessitated the placement of a left ventricular assist device (25 mL Berlin heart). He was listed for high urgency cardiac transplant and received an orthotopic heart transplant 20 days later. Hypomagnesemia was first noted at admission. Renal evaluation after cardiac transplant confirmed renal magnesium wasting, renal ultrasound revealed bilateral medullary nephrocalcinosis. Analyses of serum and urine electrolytes indicated persisting hypomagnesemia due to renal magnesium wasting accompanied by hypercalciuria. All other electrolytes as well as acid base metabolism were within normal limits.

For the evaluation of dilated cardiomyopathy, genetic testing for copy number variations (CNVs) using a microarray as well as comprehensive testing for mutations using a cardiomyopathy gene panel were performed with negative results. In addition, a genetic testing of known hypomagnesemia genes (including *CLDN16, CLDN19, CNNM2, EGF, HNF1b, and TRPM6*) failed to detect pathogenic variants.

**Family 2**

A three-year-old girl (born 1985) presented in the clinic with tetanic convulsions. She had hypocalcemia (1.1 mmol/L) and hypomagnesemia (0.5 mmol/L). During disease manifestation elevated plasma renin activity (32 ng/ml/h) and aldosterone levels (1.57 ng/ml) were noted in the absence of clinical signs of salt and/or water loss (no polyuria/polydipsia) and blood pressure was normal. Despite a treatment with calcium and vitamin D supplements, serum calcium levels did not increase until magnesium supplementation was started. Thereafter, the tetanic convulsions stopped and serum calcium concentrations normalized while serum magnesium levels remained low. Further clinical and biochemical workup clearly demonstrated a renal magnesium loss as shown in two separate parenteral magnesium loading tests (Figure 1B). In addition, when serum calcium levels were normalized, repeated episodes of hypercalciuria (6 to 12 mg/kg/d) were noted, and she developed discrete medullary nephrocalcinosis. The girl was treated with magnesium supplementation, potassium citrate, and occasionally calcium. During follow-up, the compliance with therapy was poor and serum calcium levels fell below the normal range repeatedly. At the age of nine years, the girl withdrew therapy completely.

At 12 years of age, she developed heart failure and the diagnosis of dilated cardiomyopathy was established. In addition, severe hypomagnesemia (min. 0.19 mmol/L) was noted, while serum calcium levels and urinary calcium excretions were normal. An endomyocardial biopsy was performed and did not demonstrate any abnormal finding. The mitochondrial respiratory chain activity was normal. Supportive therapy was initiated with digoxin, furosemide, captopril and fluindione. At 25 years of age, progressive heart failure necessitated external cardiac assistance. A few weeks later, she underwent successful heart transplantation. Histology of the explanted heart showed unspecific dilated cardiomyopathy.

Family analysis did not reveal further affected family members, blood and urinary magnesium levels of the parents were normal at several occasions. Renal ultrasound was normal in both parents. Mild mitral regurgitation in the father and slight left ventricular hypertrophy in the mother were detected by echography.

**Family 3**

The girl was born at 35 weeks of gestation after a pregnancy complicated by polyhydramnios with a birth weight of 2580g. During the early neonatal period, she showed muscular hypotonia. An extensive metabolic work-up, brain MRI, electromyography, as well as genetic studies for Prader-Willi syndrome were performed with unremarkable findings. At 3 weeks of age, she presented with tachycardia (heart rate of 250 beats per minute). An echocardiogram revealed left ventricular dilation as well as a ventricular septal defect of 6mm with a gradient of 36mm. Heart failure treatment with digoxin and furosemide was initiated. At 9 months of age, the diagnosis of dilated cardiomyopathy was made (table 1). After a generalized cerebral seizure episode at 2 years of age, hypokalemia and hypomagnesemia were detected and furosemide treatment was stopped. The co-existence of polyuria and polydipsia along with an impaired renal urine concentrating ability suggested a renal tubular defect. Renal ultrasound revealed medullary nephrocalcinosis. Supplementation with oral potassium and magnesium was initiated. Her cardiac function intermediately stabilized under treatment with digoxin, lisinopril, and spironolactone. An endomyocardial biopsy did not reveal signs of mitochondrial disease. Unfortunately, her cardiac function deteriorated during further follow-up, she developed cardiac failure, and finally received a cardiac transplant at 15 years of age.

Serum magnesium levels have constantly remained low (0.4 to 0.5mmol/L) despite high dose oral magnesium supplementation. Furthermore, laboratory analyses during follow-up demonstrated hypokalemia despite continuous oral supplementation as well as mild metabolic alkalosis.

After extensive analyses of known genes for renal tubular disorders including renal salt and magnesium wasting yielded negative results, whole exome sequencing was performed and analyzed together with the exomes of the individuals from families F1, F2, and F4 which finally identified the *RRAGD* gene as the single shared gene with pathogenic variants.

**Family 4**

The index case was born at 37 weeks of gestation with a weight of 2900g. There was marked polyhydramnios during pregnancy. Postnatally, polyuria was noted, but serum electrolytes remained normal. Moreover, there was no increased renin activity. At that time, there was no hypercalciuria nor nephrocalcinosis.

He presented in the clinic at 6 years of age with complaints of cramps and tetany. Laboratory findings revealed the presence of marked hypomagnesemia associated with hypokalemia. Serum calcium and sodium levels were within normal ranges.

Family analysis indicated that his father had hypokalemia, severe hypomagnesemia and nephrocalcinosis. He developed dynamic mitral valve dysfunction with severe left ventricular dysfunction. This evolved to dilated cardiomyopathy with biventricular dysfunction and he died at 41 years of age from cardiac insufficiency.

**Family 5**

After an uneventful pregnancy and birth, the girl showed feeding difficulties, emesis, and failure to thrive accompanied by paleness and sweating. An acute deterioration of her clinical status with tachycardia and tachypnea lead to admission to a local children`s hospital at 7 months of age. Under the suspicion of decompensated cardiac insufficiency, the girl was transferred to pediatric cardiology of a tertiary care hospital. Echocardiography on admission revealed a massively dilated left ventricle with a shortening fraction of <10%. Treatment for heart failure with furosemide, spironolactone, captopril, and digoxin was initiated which lead to a stabilization of cardiac function, however, the left ventricular shortening fraction remained low at 15-20%. An extensive work-up revealed no underlying metabolic abnormality or infectious cause; a cardiac biopsy was performed without decisive findings. Hypomagnesemia (serum magnesium of 0.38mmol/L) was detected at 15 months of age during a hospitalization for an airway infection. Polyuria and polydipsia were first noted during the second year of life along with additional electrolyte abnormalities including hyponatremia and hypokalemia. Urine analyses revealed hypercalciuria, ultrasound examinations in the second year of life revealed nephrocalcinosis. The girl was closely followed by a pediatric cardiologist. Fortunately, her cardiac function remained stable to date (table 1) under heart failure medication with carvedilol, enalapril, spironolactone, and hydrochlorothiazide. Digoxin treatment was stopped at 5 years of age. Over the years, she required repeated hospitalizations for airway and urinary tract infections. Polyuria as well as electrolyte abnormalities with hypomagnesemia and hypokalemia persisted despite oral supplementation. Medullary nephrocalcinosis is mild (grade 1 to 2a according to Hoyer), her renal function is actually normal.

Genetic analysis using a custom multi-gene panel for 174 cardiovascular genes (Institute for the Genetics of Cardiac Disorders, Department of Cardiology, University Hospital Münster, Germany) did not reveal any pathogenic variants in known cardiomyopathy genes.

**Family 6**

The patient presented at the age of 14 years with chest pain and NYHA Class 1 dyspnea. At that time, the echocardiography showed hyperechogenicity of the interventricular septum, with a slightly reduced ejection fraction (EF; at ~50%). The cardiologic follow-up, including stress tests and magnetic resonance imaging (MRI) following adenosine infusion, did not reveal any signs of myocardial ischemia. Blood pressure (BP) levels remained chronically low (~90/60 mmHg). Laboratory analyses showed hypomagnesemia (0.42 mmol/L) and borderline hypokalemia (3.5 mmol/L). Kidney function was preserved, with no proteinuria. The patient was treated by perindopril and bisoprolol. At the age of 35 years, the patient developed a cardiogenic shock, with severely impaired left ventricular (LV) EF (at ~15%) and systolic BP levels < 80 mmHg, in the context of pneumonia-induced sepsis. The implantation of a Boston defibrillator was necessarily performed. The patient progressively recovered to NYHA Class 2, and the latest echocardiography at the age of 36 years showed dilated cardiomyopathy (DCM) with LV EF of 45% under saccubitril/valsartan, spironolactone, bumetanide and bisoprolol. Laboratory analyses showed normal serum potassium levels, but low serum magnesium levels (0.62 mmol/L) despite oral supplementation. The father of the patient had died at the age of 33 years from myocardial infarction in a context of untreated hypercholesterolemia and poorly controlled type 2 *diabetes mellitus*. On the basis of the hetero-anamnesis of the spouse, there was no indication of DCM since the cardiac check-up performed few months before the patient died was considered as normal. The mother of the patient was asymptomatic. Her kidney function was preserved. The echocardiography showed no dilation of the cardiac cavities, with a 58% EF. The patient's grand-father and paternal uncle both died of sudden death at the age of 26 and 19 years, respectively. The autopsy of the uncle disclosed congestive DCM, with multiple intra-cardiac thromboses in a putative context of viral myocarditis. No precise clinical or biological or genetic information could be obtained about these 2 cases.

**Family 7**

The index patient was born at 38 weeks of gestation by caesarian section with a birth weight of 4250g. During pregnancy, the mother had developed gestational diabetes. The family history was remarkable for an older sister who had died in infancy during an acute febrile episode.

In infancy, she presented with recurrent episodes of vomiting, feeding problems, and failure to thrive. Lactose intolerance was diagnosed and she received lactose-free formula nutrition. At 2 years of age, she was admitted to a local hospital for acute gastroenteritis. Abdominal ultrasound revealed bilateral medullary nephrocalcinosis. Serum electrolytes were remarkable for an increased serum calcium of 2.9mmol/L, serum magnesium levels were not determined. Three months later, she was referred to a pediatric nephrology department for a diagnostic work-up of her nephrocalcinosis. At that time, hypomagnesemia (0.62 mmol/L) was detected, while all other electrolytes were within normal limits (table 1). An evaluation of mineral metabolism revealed an increased intact PTH of 90.8 pg/mL, urine analyses indicated combined renal calcium and magnesium wasting. A supplementation with oral magnesium citrate was initiated. Under the suspicion of familial hypomagnesemia with hypercalciuria and nephrocalcinosis (FHHNC; MIM #248190), genetic testing was performed which excluded mutations in *CLDN16* and *CLDN19* genes. During follow-up, feeding improved and the patient grew between the $10^{th}$ and $25^{th}$ percentile for height and weight. Nephrocalcinosis as well as hypomagnesemia and renal calcium and magnesium wasting persisted. Due to the unknown etiology of her renal phenotype, the patient was subjected to whole exome sequencing. The analysis of her single exome at that time did neither reveal pathogenic variants in known genes for hypomagnesemic disorders nor, after sequential filtering, any promising coding gene variants in homozygous or compound-heterozygous state (under the assumption of recessive inheritance).

After discovery of *RRAGD* variants in families F1 to F3, the re-analysis of her exome revealed the p.Pro119Leu variant in *RRAGD* in heterozygous state. The molecular diagnosis prompted a cardiac evaluation which revealed a dilated left ventricle with impaired pump function (FS 18%) and mitral regurge grade 1 to 2. Fortunately, a heart failure therapy with lisinopril and spironolactone resulted in a stabilization of her cardiac function and improvement of her clinical status. Since then, she is closely followed by pediatric cardiology and nephrology departments. Hypokalemia detected during a recent follow-up visit (at 5 years of age) resulted in an oral potassium supplementation in addition to magnesium. Nephrocalcinosis is stable, her renal function normal.

A re-evaluation of the autopsy report of her older sister revealed dilative cardiomyopathy with suspected acute cardiac failure in the context of a pneumonia. Unfortunately, DNA from her sister was not available. However, an analysis of the parents` DNA was initiated under the assumption of dominant inheritance. Conventional Sanger sequencing demonstrated an uncertain pathologic peak at the position of the patient`s mutation in the *RRAGD* gene in her mother. Next-generation sequencing (see above, Supplemental Methods) revealed the *RRAGD*-c.356C>T variant in 17% of analyzed reads indicating maternal mosaicism for the p.Pro119Leu mutant. An echocardiography of the mother was normal with an ejection fraction (EF) of 55%. She is treated with an ACE inhibitor for familial essential hypertension.

**Family 8**

The patient originates from a Turkish family with parental consanguinity. Two apparently healthy sisters as well as the parents do not display electrolyte abnormalities or signs of renal salt or water loss. The patient was born at term with a birth weight of 3800g. The infancy period was characterized by episodes of vomiting and fever, however, no detailed laboratory analyses were performed in local hospitals. He clinically presented at 6 years of age with a cerebral seizure. At that time, laboratory analyses revealed hypomagnesemia, hypokalemia, and hypocalcemia. In addition, metabolic alkalosis was present. Renal ultrasound was remarkable for medullary nephrocalcinosis. Potassium as well as magnesium supplementation was initiated. Genetic studies by conventional Sanger sequencing failed to detect pathogenic variants in genes for renal salt wasting (Bartter syndrome) as well as in *CLDN16* or *CLDN19* (FHHNC). Follow-up examinations demonstrated persisting hypokalemia and hypomagnesemia despite oral supplementation. Urinary analyses indicated varying calcium excretions, serum intact PTH levels were mostly within the normal range with single elevated values.

The patient is now 21 years old. His weight is 84 kg and his height 162cm (BMI of 32 $kg/m^2$). He displays stable serum potassium levels, but persisting hypomagnesemia under oral supplementation (table 1). He actually is a college student.

After excluding mutations in known genes for renal salt wasting and hypomagnesemia, he was subjected to whole exome sequencing (single exome). No promising variants were detected under the assumption of autosomal-recessive inheritance because of parental consanguinity. His exome was re-analyzed after discovery of *RRAGD* mutations in families 1 to 5. After discovery of the genetic defect in *RRAGD*, he was clinically re-evaluated including a cardiac work-up. Echocardiography demonstrated an unremarkable cardiac function and no signs of cardiomyopathy. A 24-h blood pressure monitoring (ABPM) demonstrated arterial normotension.

**Family 9**

The index patient of Family 9 (#III.3 in Supplementary Figure 1. Panel A) initially presented with muscular cramps, tetany and excessive muscular fatigability at the age of 20 years. No polyuria or polydipsia was reported. In addition, tooth erosions were noted. Blood pressure (BP) levels were in the lower range, i.e. 90/60 mmHg. Laboratory analyses showed hypomagnesemia (0.46 mmol/L), hypokalemia (3.5 mmol/L) and metabolic alkalosis ([$HCO3^-$], 35 mmol/L) (Supplementary Table 1). Kidney function was preserved, with no proteinuria. No kidney stones or nephrocalcinosis was documented. The cardiologic follow-up (until the age of 45 years) was unremarkable, with homogenous LV contractility and 78 % ejection fraction. His father (#II.9, symptomatic at the age of 20 years), as well as his paternal aunt (#II.6, symptomatic at the age of 24 years) and uncle (#II.5, symptomatic at the age of 20 years), showed a similar phenotype of hypomagnesemia-associated tetany, with no evidence for cardiac abnormalities or renal calcium deposits (Supplementary Table 1). Of note, their BP levels were also in the lower range (~90/60 mmHg).

The cousin (#III.1) of the index case presented with muscular cramps at the age of 10 years in the context of polyuria. Water restriction test showed no response to arginine-vasopressin administration. At the age of 21 years, the patient presented with a first episode of a renal colic. Follow-up (until the age of 29 years) showed recurrent formation of calcium-containing kidney stones, with no nephrocalcinosis *stricto sensu*. BP levels were in the lower range, i.e. 90/60 mmHg. Laboratory analyses showed hypomagnesemia (0.42 mmol/L), hypokalemia (3.4 mmol/L), and metabolic alkalosis ([$HCO3^-$], 28 mmol/L) (Supplementary Table 1). Kidney function was preserved, with no proteinuria. The patient was asymptomatic as far as the heart is concerned. No echocardiography has been performed thus far.

The index patient (#III.3) had 4 children (IV.1-4) with 3 different wives, of whom 2 presented with hypomagnesemia. The little girl IV.2 was born in a context of polyhydramnios that required multiple drainages. Still, she did not develop polyuria or enuresis in childhood. At the age of 16 years, she presented with tetany. BP levels were low (<90/60 mmHg), with recurrent episodes of symptomatic BP drops. Laboratory analyses showed hypomagnesemia (0.56 mmol/L), hypokalemia (3.4 mmol/L) and metabolic alkalosis ([$HCO_3^-$], 28 mmol/L) (Supplementary Table 1). Kidney function was preserved, with no proteinuria. No kidney stones or nephrocalcinosis have been documented. The patient was asymptomatic from the cardiac point of view. No echocardiography has been performed thus far. Finally, the little girl IV.4 presented with asymptomatic hypomagnesemia (0.62 mmol/L) discovered in the context of the present study at the age of 4 years. Note that serum magnesium levels were normal at the age of 2 years, similarly to patient III.6 who showed normomagnesemia at the age of three months but developed symptomatic hypomagnesemia with cramps and tetany at the age of 12 years. All members of Family 9 who were tested negative for the c.289A>C (p.Thr97Pro) *RRAGD* mutation, including II.4, II.8, II.10, III.2, IV.1 and IV.3, did not show any abnormalities in kidney or heart functions. There was no case of sudden death in Family 9.

**DETAILED STRUCTURE DISCUSSION**

All variants are found in the N-terminal G-domain and affect sequence motifs involved in nucleotide binding (Figure 1E). These motifs are highly conserved within small G-proteins and are proven to be important for high affinity nucleotide binding.[5] Ser76 is part of the G1 motif, also known as P-loop, Thr97 constitutes the G2 motif and Pro119 resides in the G3 motif (Figure 1E-G). These motifs are involved in the interaction with the phosphate moiety of the nucleotide. Ile221 resides in a sequence stretch corresponding to the G5 motif. Though the consensus of G5 is not fully conserved in RagD, the region is functionally conserved as it interacts with the base of the nucleotide (Figure 1G). The variants are expected to display low nucleotide affinity and fast nucleotide exchange. This may interfere with RagD mediated signaling in different manners. First, nucleotide free G-proteins display high affinity for GEFs. Therefore, variants with low nucleotide affinity can act as dominant negative by trapping GEFs. However, GEFs for RagD are unknown and this effect requires, that the mutation is not interfering with the interaction between GEF and G-protein per se. Second, nucleotide free G-proteins are typically instable and thus the RagD variants may be degraded. Third, due to the enhanced nucleotide exchange rate a higher fraction of the RagD variants may be GTP loaded than of wild type RagD. As consequence RagD•GTP dependent signaling may be enhanced. However, this effect requires, that the interaction between RagD-GTP and effector proteins is not perturbed. The mutations result in amino acid residues that are sterically more demanding and at least in some cases are incompatible with the fold observed in a crystal structure of human RagD (pdb entry 2q3f), where switch I is folded over the nucleotide. Such a closed conformation is typically found in GTP- or GDP-bound small G-proteins.[5] During nucleotide exchange switch I adopts an open conformation and a close and an open conformation of switch I were observed in the yeast homolog of RagD.[6-8] The open conformation would be compatible with the steric demand of the variants, but likely not with effector interaction.

Supplementary figure 1

A

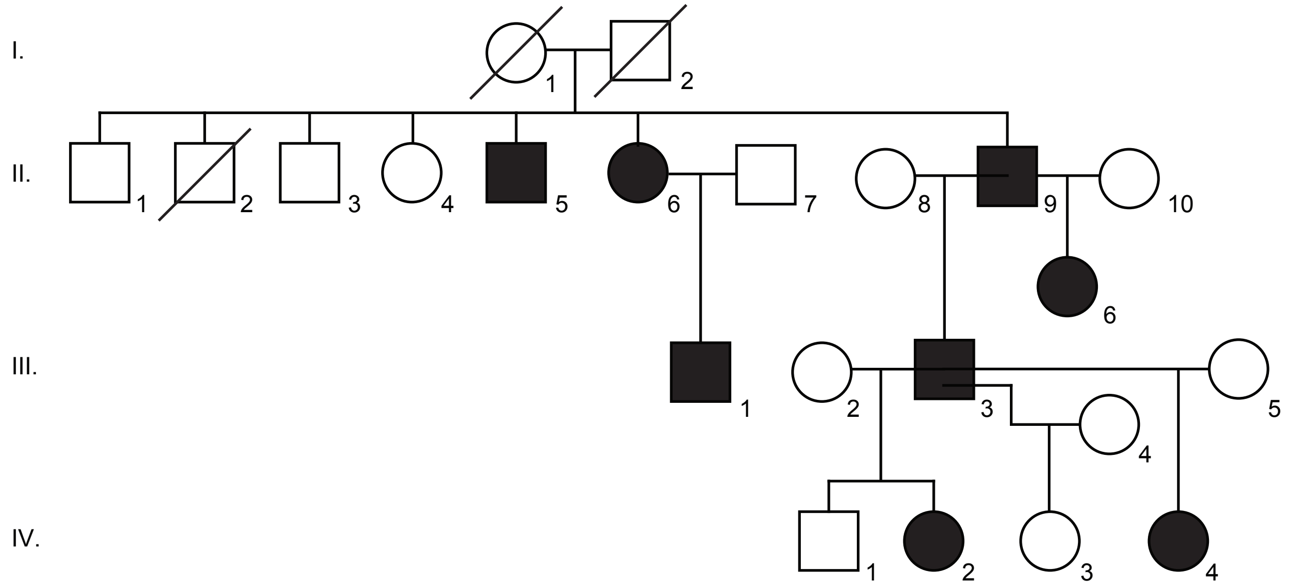

Supplementary figure 2

```
S. Cerevisiae     -----------------------------------------------------------M   1
C. Elegans        ---------------------------MESDPD---------EDFDDYRYGIDE------   18
D. Melanogaster   ----------------------------MSYDDDDYPADTFPKDFGYRAYNQDGLELEPN   32
D. Rerio          ------MTSARKNNEVDG---ENEALSFDYYDDDD-DFDAFTD--GDGDCSCDGGVLGFS   48
X. Tropicalis     -MSLPGKSH-ERQQEEEGDDEDDDIMGVSDYG----DGDSFMD--GERGSEGDDEVLDFT   52
M. Musculus       MSQVLGKPQPQGEDGGE-DEEEDELVGLAGYE----DGPESSD--AELDSGPEEGVLDFS   53
H. Sapiens        MSQVLGKPQPQDEDDAEEEEEEDELVGLADYG----DGPDSSD--ADPDSGTEEGVLDFS   54

                               G1-Box                     G2-Box
S. Cerevisiae     SLEATDSKAMVLLMGVRRCGKSSICKVVFHNMQPLDTLYLESTSNPSLEHF--STLIDLA   59
C. Elegans        EDDYPDSRPTVILMGHKRSGKTSIRKVVFQKMSPNETMFVESTARITRDTI-CSSFINFE   77
D. Melanogaster   ATGSSETKPRILLMGMRRSGKSSIQKVVFHKMSPNETLFLESTSKIVKDDINNSSFVQFQ   92
D. Rerio          DPFSSEVKPRILLMGLRRSGKSSIQKVVFHKMSPNETLFLESTNKICREDVSNSSFVSFQ   108
X. Tropicalis     DPFSTEVKPRILLMGLRRSGKSSIQKVVFHKMSPNETLFLESTNKICREDVSNSSFVNFQ   112
M. Musculus       DPFSTEVKPRILLMGLRRSGKSSIQKVVFHKMSPSETLFLESTNRICREDVSNSSFVNFQ   113
H. Sapiens        DPFSTEVKPRILLMGLRRSGKSSIQKVVFHKMSPNETLFLESTNKICREDVSNSSFVNFQ   114
                       : :  ::*** :*.**:** ****::*.* :*:::*** .   :  .  *::::.:
                   G3-Box
S. Cerevisiae     VMELPGQLNYFEPSYDSERLFKSVGALVYYVIDSQDEYINAITNLAMIIEYAYKVNPSINI   119
C. Elegans        TIEFPGQMCPFDDSLDPVGVFQKCEALLFIIDAQAELQEPIATLVEYFCRAYKINQNIKF   137
D. Melanogaster   IWDFPGQIDFFEPTFDSDMIFGGCGALVFVIDAKDDYNEALTKFKNTVLQAYKVNKRIKF   152
D. Rerio          IWDFPGQIDFFDPTFDYEMIFRGTGALIFVIDSQDDYVEALSRLHLTVTRAYKVNPDINF   168
X. Tropicalis     IWDFPGQIDFFDPTFDYEMIFRGTGALIFVIDSQDDYMEALARLHLTVTSAYKVNPDINF   172
M. Musculus       IWDFPGQIDFFDPTFDYEMIFRGTGALIFVIDSQDDYMEALARLHLTVTRAYKVNTDINF   173
H. Sapiens        IWDFPGQIDFFDPTFDYEMIFRGTGALIFVIDSQDDYMEALARLHLTVTRAYKVNTDINF   174
                    ::***:  *: :  *   :*    **:::**:: :  : :: :   .  ***:*  *::
                      G4-Box                                  G5-Box
S. Cerevisiae     EVLIHKVDGLSEDFKVDAQRDIMQRTGEELLELGLDGVQVSFYLTSIFDHSIYEAFSRIV   179
C. Elegans        EVFVHKADGLTEEARVETKFNIYHQVKETIKDQIDVDLQVTYHLTSIYDHSIFEAFSKVV   197
D. Melanogaster   EVFIHKVDGISDDSKMESQRDIHQRSSDDLNEAGLDQIHLSFHLTSIYDHSIFEAFSKVV   212
D. Rerio          EVFIHKVDGLSDDHKIEKQRDIHKRANDDLADAGLERIHLSFYLTSIYDHSIFEAFSKVV   228
X. Tropicalis     EVFIHKVDGLSDDHKIETQRDIHQRANDDLVDAGLEKIHLSFYLTSIYDHSIFEAFSKVV   232
M. Musculus       EVFIHKVDGLSDDHKIETQRDIHQRANDDLADAGLEKIHLSFYLTSIYDHSIFEAFSKVV   233
H. Sapiens        EVFIHKVDGLSDDHKIETQRDIHQRANDDLADAGLEKIHLSFYLTSIYDHSIFEAFSKVV   234
                  **::**.**:::: ::: : :* ::  : : :     ::::::****:****:****::*

S. Cerevisiae     QKLIPELSFLENMLDNLIQHSKIEKAFLFDVNSKIYVSTDSNPVDIQMYEVCSEFIDVTI   239
C. Elegans        QNLVKQLPTLERLLDVFNNSSKVTKSFLFDILSKIYIATDSEPVEMSIYELCCDMIDVTL   257
D. Melanogaster   QKLIPQLPTLENLLNIFIPNSGIEKAFLFDVVSKIYIATDSSPVDMQTYELCCDMIDVVI   272
D. Rerio          QKLIPQLPTLENLLNIFISNSGIEKAFLFDVVSKIYIATDSSPVDMQTYELCCDMIDVVI   288
X. Tropicalis     QKLIPQLPTLENLLNIFISNSGIEKVFLFDVVSKIYIATDSSPVDMQTYELCCDMIDVVI   292
M. Musculus       QKLIPQLPTLENLLNIFISNSGIEKAFLFDVVSKIYIATDSTPVDMQTYELCCDMIDVVI   293
H. Sapiens        QKLIPQLPTLENLLNIFISNSGIEKAFLFDVVSKIYIATDSTPVDMQTYELCCDMIDVVI   294
                  *:*: :*  **.:*: :   * : * ****: ****::*** **::. **:*.::***.:

S. Cerevisiae     DLFDLYKAPVLRNSQKSSDKDNVINPRNELQNVSQLANGVIIYLRQMIRGLALVAIIRPN   299
C. Elegans        DLSSIYGVAEN--G---------SNYDERSSSVIRLKSEQVMFLRQVNKHLALVFIMKED   306
D. Melanogaster   DLSSIYSSE----E---------TAFDSGSSSLIKLNNNTILYLREVNKFLALVCILREE   319
D. Rerio          DISCIYGLSGDEGG---------TPYDKESMAIIHLNNTTVMYLKEVTKFLALVCFLREE   339
X. Tropicalis     DISCIYGLE--GAG---------TPYDKESLAIIKLNNTTVLYLKEVTKFLALVCFVREE   341
M. Musculus       DISCIYGLKEDGAG---------APYDKDSTAIIKLNNTTVLYLKEVTKFLALVCFVREE   344
H. Sapiens        DISCIYGLKEDGAG---------TPYDKESTAIIKLNNTTVLYLKEVTKFLALVCFVREE   345
                  *:  :*                         .    : :* .  :::*::: : **** ::: :

S. Cerevisiae     GTDMESCLTVADYNIDIFKKGLEDIWANARASQAKNSIEDDV------------------   341
C. Elegans        GNEKA---GFIDHNFGVFKAGIEQVFKVKNRGVNF-------------------------   338
D. Melanogaster   NFNRQ---GVIDYNFICFRDAISEVFELRLKRQKQLENNDQDDDLVDEQTLIRHGHDAG   376
D. Rerio          SFEKK---GLIDYNFHCFKKAIEEVFDVRLKVQRSRKLLSQRRWSR----QTVPN-----   387
X. Tropicalis     SFERK---GLIDYNFHCFRKAIQEVFEVRVKVLRSRKHQSQTKKSR----RATPN-----   389
M. Musculus       SFERK---GLIDYNFHCFRKAIHEVFEVRMKMVKSRKAQSRLPKKT----GATPN-----   392
H. Sapiens        SFERK---GLIDYNFHCFRKAIHEVFEVRMKVVKSRKVQNRLQKKK----RATPN-----   393
                  . :      . *:*:  *: .: :::

S. Cerevisiae     ---------       341
C. Elegans        ---------       338
D. Melanogaster   GISRAQPIN       385
D. Rerio          GTQV-LPH-       394
X. Tropicalis     GTPG-VPL-       396
M. Musculus       GTPR-VLL-       399
H. Sapiens        GTPR-VLL-       400
```

Supplementary figure 3

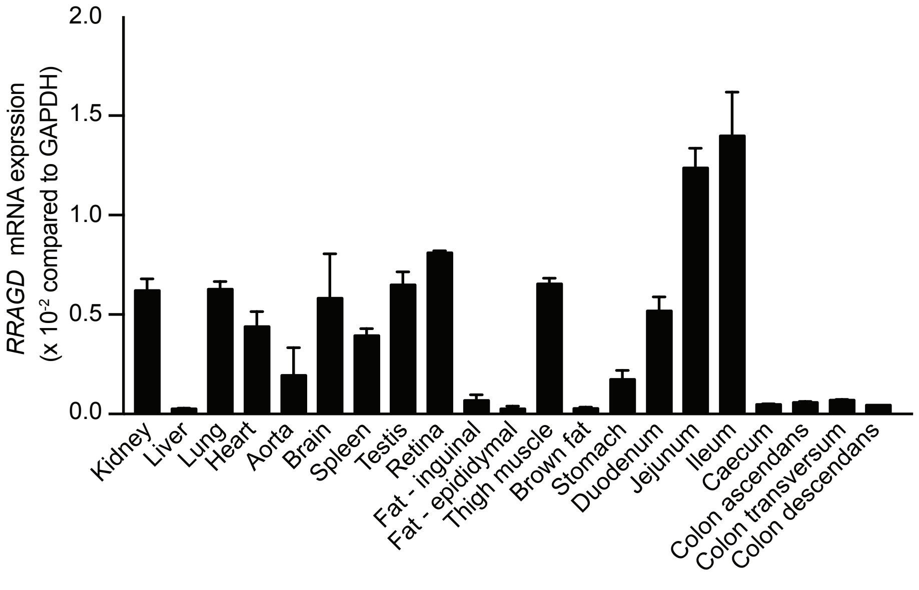

Supplementary figure 4

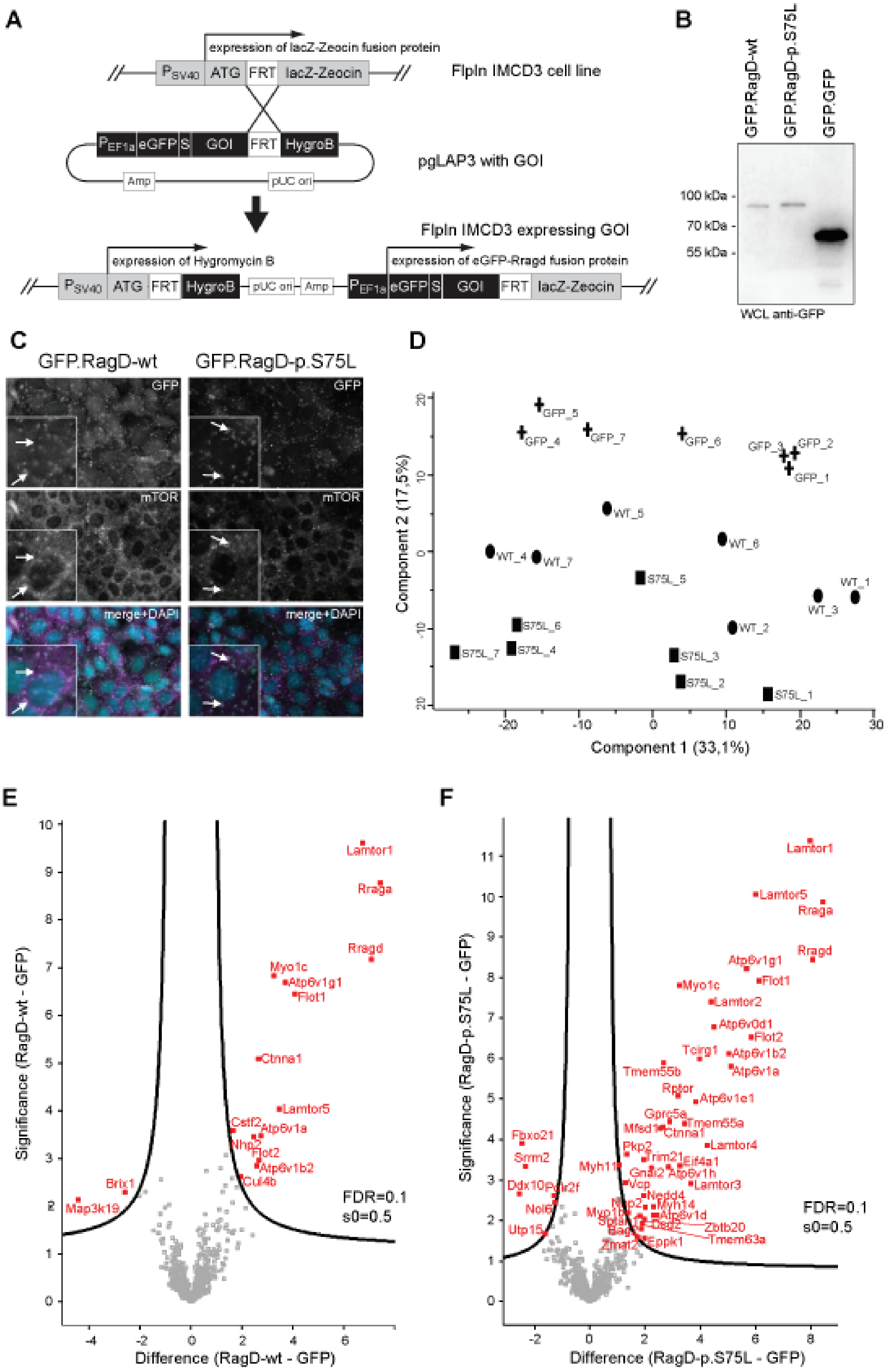
